# Myosin-driven actin-microtubule networks exhibit self-organized contractile dynamics

**DOI:** 10.1101/2020.06.11.146662

**Authors:** Gloria Lee, Michael J. Rust, Moumita Das, Ryan J. McGorty, Jennifer L. Ross, Rae M. Robertson-Anderson

## Abstract

The cytoskeleton is a dynamic network of proteins, including actin, microtubules, and myosin, that enables essential cellular processes such as motility, division, mechanosensing, and growth. While actomyosin networks are extensively studied, how interactions between actin and microtubules, ubiquitous in the cytoskeleton, influence actomyosin activity remains an open question. Here, we create a network of co-entangled actin and microtubules driven by myosin II. We combine dynamic differential microscopy, particle image velocimetry and particle-tracking to show that both actin and microtubules in the network undergo ballistic contraction with surprisingly indistinguishable characteristics. This controlled contractility is distinct from the faster turbulent motion and rupturing that active actin networks exhibit. Our results suggest that microtubules can enable self-organized myosin-driven contraction by providing flexural rigidity and enhanced connectivity to actin networks. These results provide important new insight into the diverse interactions cells can use to tune activity, and offer a powerful platform for designing multifunctional materials with well-regulated activity.

## Introduction

The cytoskeleton is a dynamic composite network of proteins that not only provides cells with mechanical support but also restructures dramatically to enable essential biological processes such as cell division, growth, mobility, and intracellular transport^1–3^. These diverse dynamics and properties are enabled by the inherent activity of the actin filaments, microtubules, and motor proteins that comprise the cytoskeleton. One ubiquitous interaction the cytoskeleton uses to self-organize and produce active work is the association of the motor protein myosin II with actin filaments. These actomyosin complexes generate contractile motion and force by myosin II minifilaments pulling actin filaments past each other to drive cell migration, signaling, adhesion, protrusion, and retraction^4–7^.

A critical parameter that influences the large-scale activity of actomyosin networks is the connectivity of the network, which can be tuned via actin crosslinking proteins^8–10^. In vitro studies have shown that actin networks must be crosslinked above a critical percolation threshold to sustain ordered contractility across the network^8–11^. Without additional crosslinkers linking actin filaments together, actomyosin activity in disordered actin networks can induce both contractile and extensile forces, resulting in flow and rupturing of the actin network^12,13^. In vitro studies have shown that in this regime myosin restructures actin into disconnected foci and the system exhibits turbulent motion rather than organized contraction^14^. Additional crosslinkers allow myosin motors to generate internal stress more efficiently in the network as they mobilize actin^15–17^. In response to this stress, actin filaments can buckle, shorten, or even break, leading to self-organized network contraction^18^.

Microtubules can also undergo active rearrangement via interactions with motor proteins such as kinesin^19–21^. Studies on kinesin-driven microtubule networks have shown that the kinesin-microtubule interaction generally lead to extensile rather than contractile motion and often results in flow rather than coarsening or restructuring^20–23^, though microtubule contraction has been observed as well^19,24^.

While active actin networks and microtubule networks have been intensely studied over the last decade^4,25^, how the presence of one filament type affects the active dynamics of the other remains an open question^26^. At the same time, interactions between actin and microtubules are pervasive in the cell and are particularly important for regulating cell shape during migration, division, and intracellular transport^27^. Thus, understanding how the composite nature of the cytoskeleton affects its motor-driven activity is of critical importance. Further, we have previously shown that in vitro systems of co-entangled actin and microtubules exhibit novel mechanical properties, absent in single-component networks, that could be harnessed for the design of multifunctional materials^28–30^. As such, incorporating activity into this system could provide a powerful platform for engineering active non-equilibrium materials.

Here, we create, for the first time to our knowledge, an active composite network of co-entangled actin and microtubules that exhibits contractility driven by myosin II minifilaments. We use two-color confocal fluorescence microscopy to visualize the network reorganization during motor activity. We use particle image velocimetry and dynamic differential microscopy, complemented by particle-tracking experiments, to characterize the active network dynamics. We show that microtubules enable organized and uniform contraction of actomyosin networks that otherwise exhibit disjointed and turbulent contraction dynamics. Actin-microtubule networks also exhibit slower and more sustained contraction than actomyosin alone, while still supporting nearly ballistic contractility characterized by steady directed motion. Further, our particle-tracking experiments show that particles embedded in active actin-microtubule networks exhibit two distinct transport modes dependent on the timescale probed. At small timescales, particles move subdiffusively, similar to in steady-state networks, while at larger timescales the motion is ballistic, akin to actomyosin networks. Our findings suggest that actin-microtubule interactions may play an important role in tuning actomyosin activity in cells, and could be exploited in cytoskeleton-inspired active matter to not only tune the stiffness of the material but also the spatial and temporal scales of activity.

## Results

### Designing an active actin-microtubule network

We designed active actin-microtubule networks using myosin II to drive activity (SI Videos 1-3). While active actin networks and microtubule networks have been intensely investigated in recent years^4,25^, to our knowledge, active matter comprised of both actin filaments and microtubules have yet to be realized. We previously determined optimal buffers and polymerization conditions to co-polymerize actin and microtubules to form composite networks that are homogeneous and in which the two filament types are co-entangled with one another^28–30^. Here, we incorporate myosin II activity into these networks to bring this system one step closer to the cellular cytoskeleton, and to provide new design principles for active biomaterials. We focus on networks with equal molarity of actin and tubulin, since we have previously shown that networks with this actin to tubulin ratio exhibit strong resistance to imposed stress while maintaining a high degree of filament mobility^29^. This combination of resilience and mobility provides an ideal platform for designing active materials that can respond to stimuli while maintaining structure.

To produce activity, we incorporate myosin II minifilaments at a 1:12 myosin to actin molar ratio, which we determined to produce the highest degree of activity that did not result in network destruction or irreproducible activity. To control activity, we incorporate a saturating concentration of blebbistatin into the networks which de-activates myosin II activity except when exposed to ~400-500 nm light^31^. We use two-color fluorescence confocal microscopy to image the actin and microtubule networks and initiate activity. We use 488 nm light to simultaneously visualize the actin channel and activate myosin-driven activity, and we use 561 nm light to image microtubules. To our knowledge, this is the first active network of co-entangled actin and microtubules synthesized *in vitro*. As we describe below, this three-dimensional system exhibits robust and uniform myosin-driven contractility.

### Actin-microtubule contraction is synchronized and organized

Figure 1A shows images from a representative time-series of an actin-microtubule network undergoing myosin-driven activity over the course of six minutes. The top row displays dual-color images with both the actin and microtubule channels visualized, while the bottom two rows show the separate actin and microtubule channels. From the images, it is clear that myosin activity causes the network to move and rearrange. We see more clustering at later time points, as well as conformational changes and mobility of some of the brighter structures (indicated by arrows). Specifically, the horizontal set of arrows shows co-localized actin filaments and microtubules disappearing from the plane of view over time. The vertical set of arrows shows co-localized actin filaments and microtubules changing shape in the plane of view. Surprisingly, despite the fact that myosin only binds to actin, the two networks remain well-integrated and microtubules co-localize and move with the actin filaments. While we only analyze the contraction dynamics for 6 minutes, we see the well-incorporated composite network contract for up to 30 minutes.

**Figure 1.**
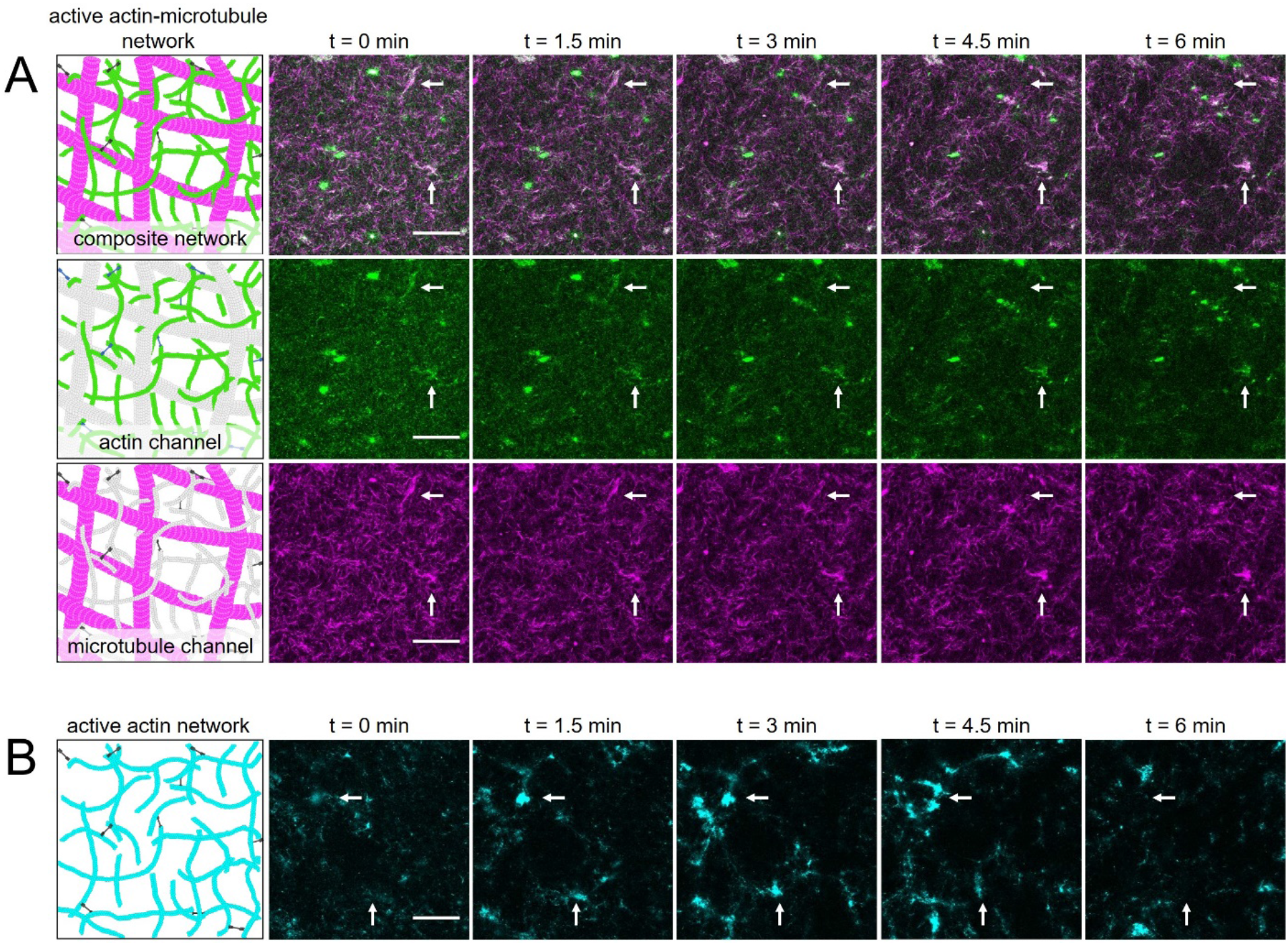
Design and characterization of active actin-microtubule networks. (A) (Column 1) Cartoon depictions of an active actin-microtubule network comprised of actin filaments (green/grey), microtubules (magenta/grey) and myosin minifilaments (black). As depicted, the actin filaments and microtubules are co-entangled with one another and the myosin can bind to actin to generate activity. (Columns 2-6) 256×256 pixel confocal microscopy images of an active actin-microtubule network at five equally spaced time points of a six minute time-series (SI Video 1). Two-color images in the top row display both actin (green) and microtubules (magenta) in the composite network. The second and third rows separately show the actin (SI Video 2) and microtubule (SI Video 3) channels, respectively. Either actin or microtubules are shaded grey in the corresponding cartoons to reflect the imaging channel not shown. Arrows in the images point to examples of co-localized actin filaments and microtubules moving and reconfiguring due to myosin activity. Scale bar is 50 μm. (B) Cartoon depiction and corresponding confocal micrographs of an active actin network (SI Video 4). Actin filaments are colored cyan to differentiate from the actin channel in the actin-microtubule networks. The network is prepared identically to the active actin-microtubule network but lacks microtubules. Arrows in the images point to examples of actin moving and reconfiguring due to myosin activity. Scale bar is 50 μm.

As a control, to determine the effect of actin-microtubule interactions on actomyosin activity, we created active networks with the tubulin removed but with all other reagents and conditions fixed (SI Video 4). As shown in Figure 1B, these active actin networks undergo substantially more restructuring and rupturing during myosin activity, as actin filaments form bright bundles and foci and leave large voids in the network. The horizontal set of arrows shows a large bundle of actin forming then disappearing from the plane of view, while the vertical set of arrows indicates a region where a bundle of actin enters then exits the plane of view. As such, microtubules appear to provide stabilization to actomyosin networks, and enable slower, more controlled actomyosin activity. Static actin-microtubule networks, either without light-activation or without myosin, showed no network rearrangement over the six minutes of imaging (SI Videos 5-6).

To determine the direction of motion of the filaments in the active networks, we used Particle Image Velocimetry (PIV) analysis to generate velocity vector fields at different times during activity. Figure 2A shows the velocity fields for the actin (top row) and microtubule (bottom row) channels of an active actin-microtubule network. At each time point and for each channel, the velocity vectors point towards the center of the activated region, showing that the network is contracting over time. Importantly, the velocity fields for actin and microtubules are quite similar at each time point, demonstrating that the microtubules and actin move together. Further, the velocity fields for all time points are similar, pointing inward with similar magnitudes, indicating that the network is undergoing controlled and organized contraction.

**Figure 2.**
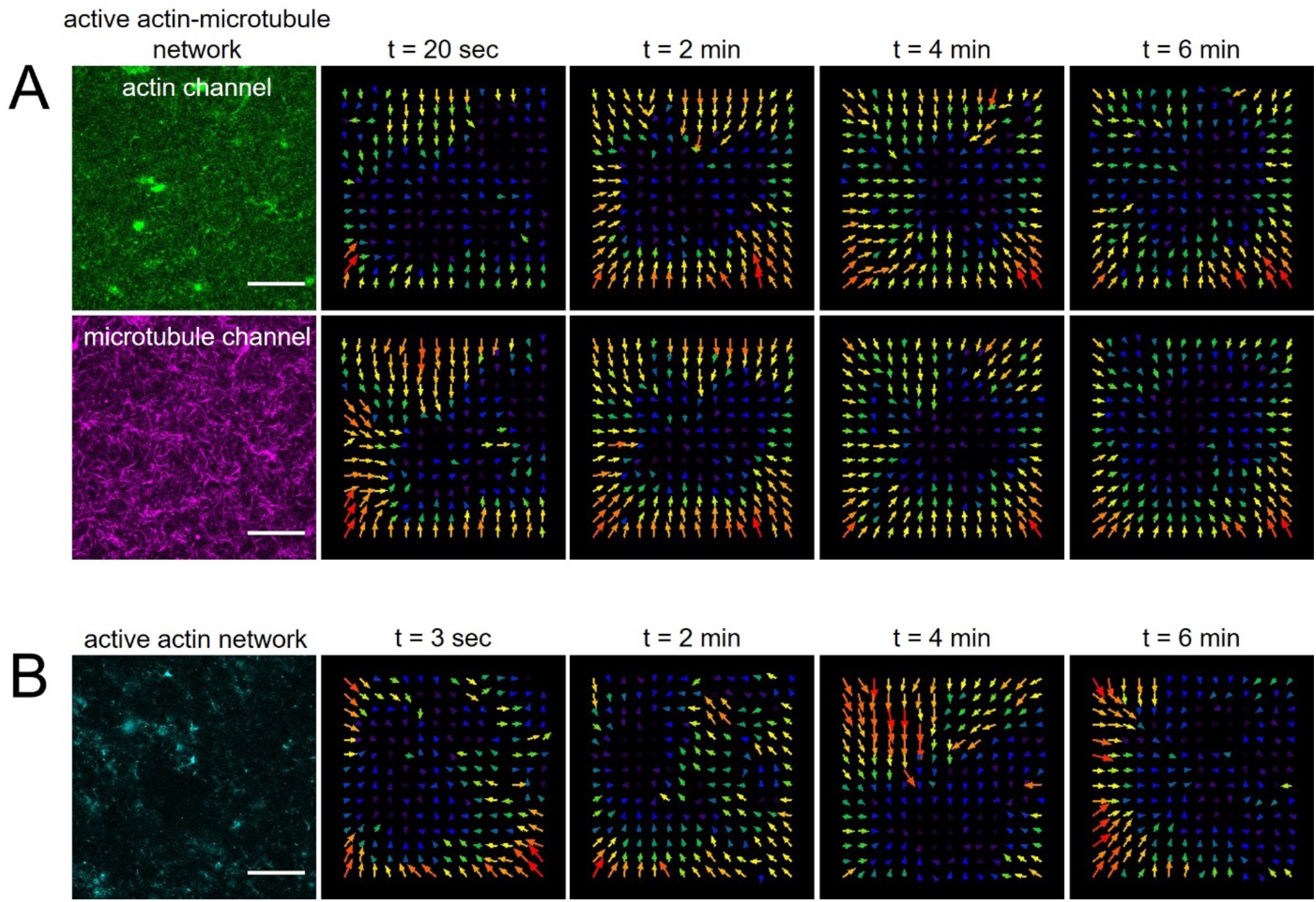
Particle Image Velocimetry shows that microtubules enable organized myosin-driven contraction. (A) Velocity vector fields of the actin channel (green, top row) and microtubule channel (magenta, middle row) of the active actin-microtubule network shown in Figure 1 at four time points during the 6 minute time-series. Velocity fields were generated using PIV analysis with a lag time of 20 s. Velocity fields for both channels show consistent motion directed towards the center region of the field-of-view throughout video duration. Scale bars in all images are 50 μm. (B) Velocity vector fields of the active actin network (cyan), generated via PIV analysis with a lag time of 3 s, show faster and more turbulent motion than the actin-microtubule network throughout the duration of the time-series. Scale bars in all images are 50 μm.

Unlike the composite network, the active actin network, shown in Figure 2B, exhibits velocity fields that quickly change direction and magnitude over time. Moreover, while the velocity fields for the composite actin-microtubule networks were computed using frames that were 20 s apart, the actin networks needed frames that were only 3 s apart. When we tried to use longer lag times, the velocity fields were dominated by noise and important features of the network escaped the field of view. This result implied that the contraction speed was faster and less controlled in the actin network compared to the composite. While the composite network exhibits characteristics of controlled network-wide contraction, the actin network ruptured and contracted towards different foci that moved in a turbulent fashion over time, as previously reported for actomyosin networks without additional crosslinkers^12,13^. This difference suggests that microtubules can organize actomyosin activity as well as slow it down.

### Microtubules slow actomyosin activity 7-fold

To quantify the active dynamics of myosin-driven networks, we use differential dynamic microscopy (DDM) to analyze time-series of images. DDM is useful for investigating diffusive and active dynamics of soft materials such as those found in biological systems^32^, including cytoskeleton complexes^33–35^. In our own previous work, we have used DDM to study the passive transport of tracers within static actin-microtubule networks, and found that these networks exhibit subdiffusive behavior^36^. Here, we first use DDM analysis to generate image structure functions *D*(*q*, *Δt*) at varying wavenumbers *q*, as described in the Methods. Figure 3A shows *D*(*q*, *Δt*) curves for an active actin-microtubule network, increasing with lag time for three representative wavenumbers. At each wavenumber, *D*(*q*, *Δt*) curves are largely indistinguishable between the actin and microtubule channels, demonstrating the correlated motion of the two filaments. As lag time increases, *D*(*q*, *Δt*) increases and then begins to plateau, at which point density fluctuations in the network are considered no longer correlated. By fitting *D*(*q*, *Δt*) with an exponential function, as described in the Methods, we extract a characteristic decay time *τ* for fluctuations of a given wavenumber (Fig 3C). This time roughly corresponds to the lag time at which *D*(*q*, *Δt*) reaches a plateau. As shown, both actin and microtubule channels appear to approach this plateau at similar lag times. However, *D*(*q*, *Δt*) for active actin networks reach plateaus more quickly than active actin-microtubule networks, indicating faster dynamics (Fig 3B). In contrast to the active networks, *D*(*q*, *Δt*) for inactive actin-microtubule networks exhibit no plateau and can thus not be fit to extract a decay time (SI Figure 1). This lack of a plateau indicates that the dynamics of the inactive networks are much slower than active networks, and do not rearrange appreciably over the experimental time frame.

**Figure 3.**
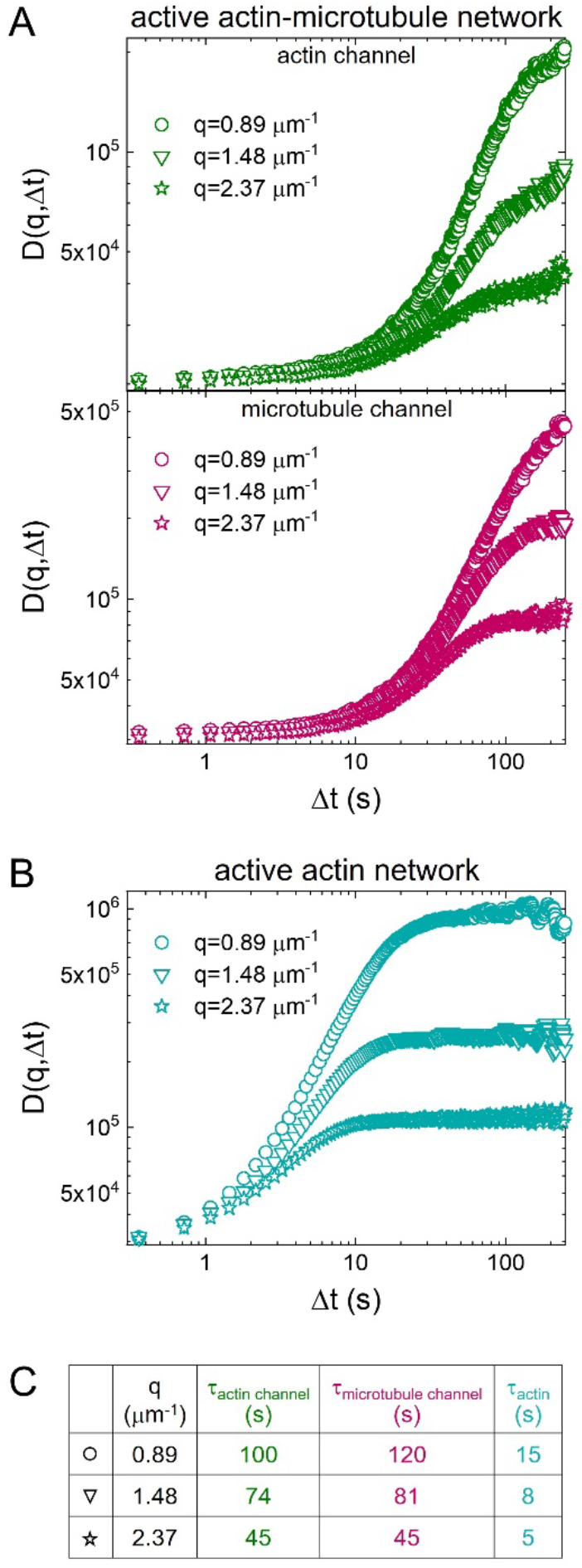
DDM demonstrates that actin and microtubules in active actin-microtubule networks exhibit nearly indistinguishable contraction dynamics that are slower than in actin networks. (A) Representative image structure functions *D*(*q*, *Δt*) for the actin channel (green, top) and microtubule channel (magenta middle) of an active actin-microtubule network, for wavenumbers of *q*= 0.89 μm^-1^ (circles, highest curves), 1.48 μm^-1^ (triangles, middle curves) and 2.37 μm^-1^ (stars, lowest curves). Note the similarity between curves for the actin and microtubule channels. (B) Representative image structure functions for the active actin network, for the same three wavenumbers *q* shown in (A). Note that *D*(*q*, *Δt*) curves for the active actin network approach plateaus at shorter lag times than for the actin-microtubule network and, as such, have shorter decay times. (C) Decay times *τ* for all curves plotted in (A) and (B), determined via fitting each curve to an exponential function (see Methods). Decay times roughly correlate with when *D*(*q*, *Δt*) curves reach a plateau and decrease as wavenumber increases. Decay times for the actin and microtubule channels of the actin-microtubule network are in close agreement, and an order of magnitude longer than those for the actin network.

To more fully capture the dynamics of the active networks we evaluate the dependence of the decay time *τ* on the wavenumber *q*. Figure 4A shows the average *τ*(*q*) curves for both the actin and microtubules of the active composite network as well as for the actin network. The curves for both actin and microtubules of the composite are nearly identical, signifying that the dynamics of both types of filaments are highly correlated. Comparing the actin network to the composite, we see that for any given wave number, the characteristic decay time of the actin network is significantly smaller than that of the actin-microtubule network, showing that the density fluctuations decay faster. This result supports our PIV analysis that shows that the actin networks exhibit much faster dynamics compared to actin-microtubule networks.

**Figure 4.**
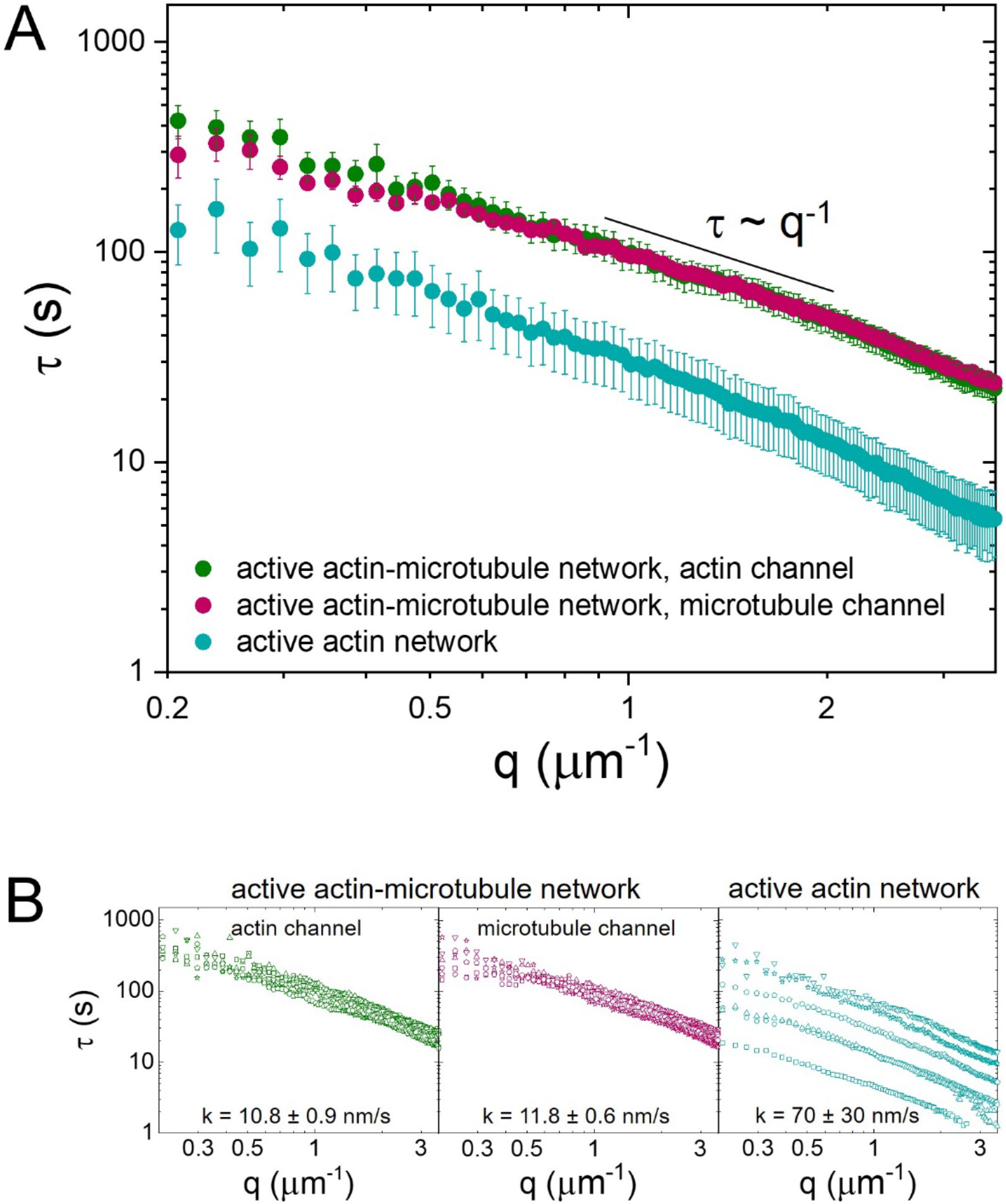
DDM reveals that microtubules slow down and organize ballistic motion of myosin-driven cytoskeleton networks. (A) Average characteristic decay time *τ* vs wavenumber *q* for the actin channel (green) and microtubule channel (magenta) of an active actin-microtubule network as well as an active actin network (teal). All curves appear to roughly follow the scaling *τ* ~ *q*^−1^ for larger *q* values, indicating ballistic motion. Smaller *τ*(*q*) values for all wavenumbers show that the active actin network exhibits faster motion than the actin-microtubule network. (B) Individual *τ* vs *q* curves from which the average curves displayed in (A) were computed. Fitting each curve to *τ*~1/*kq* provides an effective speed *k* for each filament and each network, which is listed in the corresponding plots. The large spread in active actin data compared to the active actin-microtubule data shows that the microtubules are able to organize actomyosin activity and enable controlled contraction.

The shape of the *τ*(*q*) functions reveals the type of motion the active networks exhibit and the corresponding rate of motion. If *τ* ~1/*kq*, then the system exhibits ballistic motion and *k* represents the speed. If *τ* ~ 1/*kq*^2^, then the system exhibits diffusive motion and *k* represents the diffusion coefficient. All curves for actin and microtubules in composite or actin networks exhibit roughly *q*^−1^ dependence, signifying ballistic motion. We find exponents of −1.15 ± 0.03 and −0.85 ± 0.05 for the actin channel and microtubule channel of the composite network, respectively. An exponent of −1.00 ± 0.03 is found for the active actin network.

We fixed the scaling exponent to −1 to determine an effective speed *k* from the relation *τ* ~1/*kq*. We compute contraction speeds of 10.8 ± 0.9 nm/s and 11.8 ± 0.6 nm/s for the actin and microtubules in the composite network, compared to 70 ± 30 nm/s for the active actin network (Fig. 4B). Note that the speeds for both actin and microtubules in the composite network are identical, within error, while the motion of the actomyosin network is ~7x faster. Further, the large standard error for the actomyosin network speed suggests a wide variation in activity.

Each average curve displayed in Figure 4A is computed from six different regions of interest in two different networks, displayed in Figure 4B. The curves for both channels of the actin-microtubule network are overlapping with very little spread in the data, suggesting that the contraction dynamics are highly controlled and reproducible. Conversely, the active actin network exhibits a wider spread in τ vs q curves, demonstrating the highly variable activity that actomyosin networks display in the absence of microtubules.

### Particles exhibit both subdiffusion and ballistic transport in active actin-microtubule networks

To provide another independent measure of dynamics to complement our DDM results, we perform particle-tracking analysis as described in the Methods. We compute mean-squared displacements (MSDs) of beads embedded in each network as a function of lag time *Δt*. Particles undergoing normal Brownian diffusion exhibit a scaling of *MSD*~*Δt*^1^, whereas ballistic motion manifests as *MSD*~*Δt*^2^ and subdiffusion manifests as *MSD*~*Δt*^*α*^ with *α* < 1. To determine the extent to which the network dynamics deviate from Brownian diffusion, we plot *MSD*/*Δt* vs *Δt* curves on a log-log scale (Fig 5). Subdiffusive MSDs have a negative slope and superdiffusive or ballistic MSDs have a positive slope. As expected, particles in the inactive actin-microtubule network exhibit purely subdiffusive behavior for all experimental lag times, with *MSD*~*Δt*^0.6^. This exponent is similar to that reported previously for similar actin-microtubule networks^36^. Particles in the active actin network exhibit subdiffusive behavior for the first ~2-3 s after which the motion is nearly ballistic (*MSD*~*Δt*^1.7^. Interestingly, particle dynamics in the active actin-microtubule network appear similar to the inactive network for the first ~30 s, with *MSD*~*Δt*^0.7^, after which it transitions to nearly ballistic behavior similar to the active actin network, with *MSD*~*Δt*^1.55^. This result demonstrates that thermally driven network fluctuations dictate the mechanics of the active composite network for shorter timescales, while motor-driven contraction controls the motion at larger timescales. This separability of dynamics could play important roles in allowing how the cytoskeleton responds to mechanical and chemical stimuli.

**Figure 5.**
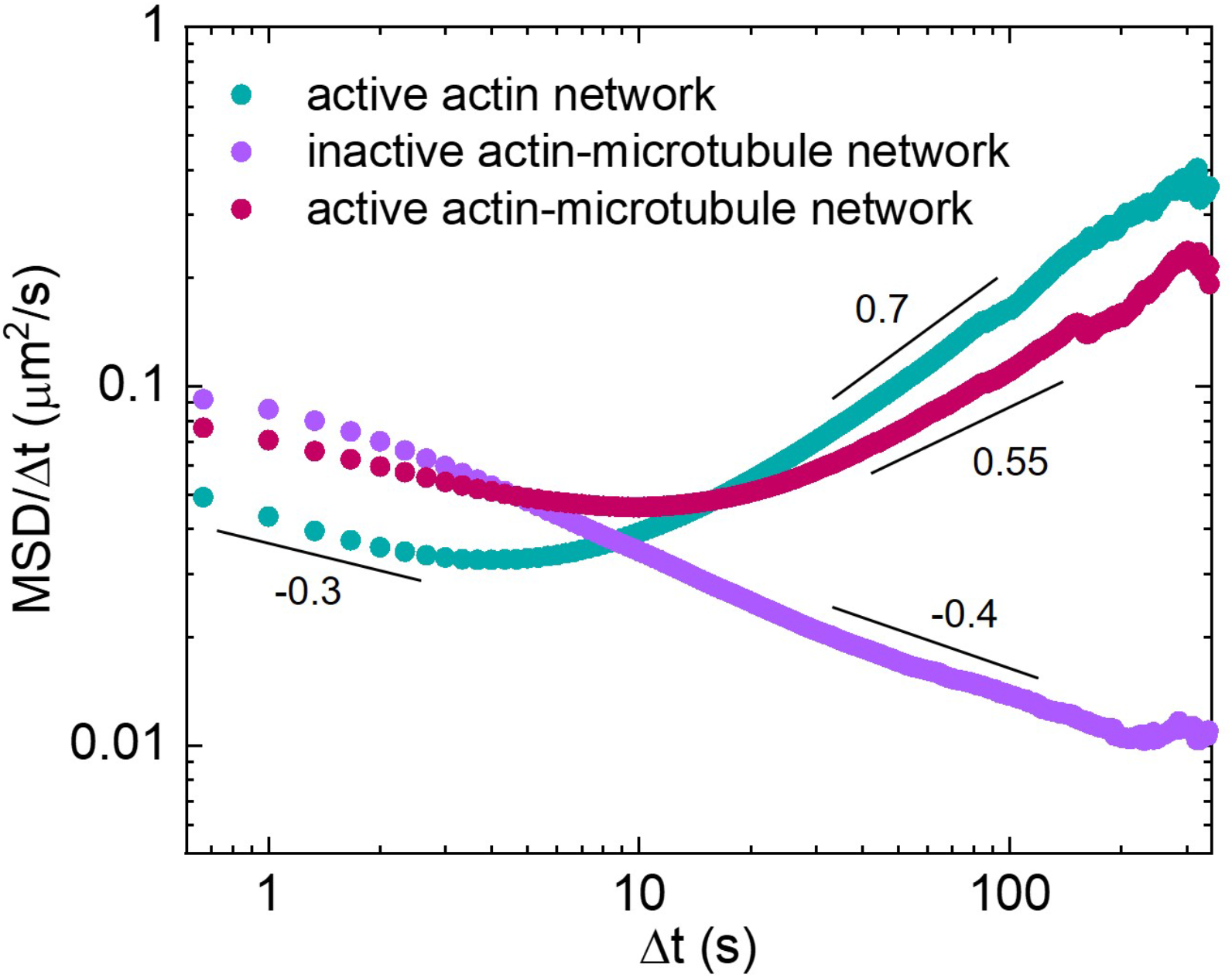
Particles in active actin-microtubule networks exhibit subdiffusive behavior at short lag times while near-ballistic motor-driven behavior dominates their long-time dynamics. Mean-squared displacements (MSD) of microspheres embedded in cytoskeleton networks, determined via particle-tracking algorithms, divided by the corresponding lag time (Δt) as a function of Δt. MSD/Δt curves for active actin-microtubule (magenta), active actin (teal), and inactive actin-microtubule (purple) networks are shown. Scaling bars and corresponding numbers show approximate power-law scaling of MSD/Δt curves. Particles in inactive actin-microtubule networks exhibit subdiffusive behavior over the entire experimental window while in active actin networks they exhibit near ballistic dynamics over the majority of the experimental lag times. In active actin-microtubule networks, on the other hand, particles exhibit subdiffusion similar to inactive networks for short lag times but transition to dynamics similar to active actin networks at large lag times.

## Discussion

Our collective results demonstrate that microtubules play an essential role in regulating actomyosin dynamics. Specifically, co-entangled actin-microtubule networks driven by myosin II exhibit organized and controlled contraction dynamics. In contrast, similar actomyosin networks that lack microtubules undergo fast, multidirectional motion and network rupturing. Importantly, while previous studies have shown that in actomyosin networks, actin crosslinking above the percolation threshold is required for organized contraction dynamics, we show here that microtubules can also enable self-organized contractile activity.

There are several ways microtubules may modulate actomyosin contraction. Microtubules have a ~100*x* greater persistence length than actin filaments, so a network of microtubules entangled within an actomyosin network may mechanically couple the composite network over larger length scales. Such connectivity is a necessary condition for actomyosin contraction^8^. In this manner, microtubules replicate the function of crosslinkers in biasing network activity away from extension towards contraction, by increasing network connectivity and facilitating propagation of local myosin activity across a larger range of the network. Previous studies have also shown that in actin-microtubule composites, actin filaments provide lateral reinforcement to microtubules, which are then able to bear larger compressive loads and stabilize the network^37^. This interplay may explain why the presence of microtubules both organizes actomyosin activity and slows the rate of network rearrangement.

Finally, previous studies have shown that filament rigidity plays an important role in determining whether motor-driven activity is contractile or extensile and the dimensionality of the dynamics^9^. Specifically, increasing filament rigidity can reduce the dimensionality of the contraction and also allow for extensile motion at low connectivity and contraction at high connectivity. As such, the rigidity of the microtubules itself could serve to reduce the degrees of freedom of the dynamics, leading to more organized and controlled motion, while, at the same time, offering extensile feedback during contraction to slow the rate of contraction. This interplay between rigidity and connectivity in tuning the active dynamics of myosin-driven networks suggests that a composite network of both rigid microtubules and semiflexible actin filaments can confer contractility and activity that cannot be reproduced in single-filament active systems.

In the cell, the cytoskeleton is a non-equilibrium composite network that self-organizes and restructures to enable processes as diverse as cell motility, mechanosensing and intracellular transport^4,8^. Many of these processes rely on interactions between actin and microtubules^6,26,38,39^ as well as active rearrangement and force generation via motor proteins such as myosin II^4^. Here, we present the synthesis and characterization of an active composite network of actin and microtubules with activity driven by myosin motors. We couple particle image velocimetry with differential dynamic microscopy to show that myosin-driven actin-microtubule networks exhibit robust ballistic contractility that is more organized and slower than networks that lack microtubules. Further, while myosin only associates with actin, the dynamics of the actin filaments and microtubules in the network are nearly indistinguishable, suggesting that they are strongly coupled and that the presence of the microtubules can serve to modulate actomyosin activity. These findings indicate that in cells, actomyosin contraction may be regulated via microtubules, in addition to chemical crosslinkers. Our particle-tracking analysis also shows that active actin-microtubule networks exhibit subdiffusive behavior at short lag times, similar to inactive networks, and near-ballistic behavior, similar to actomyosin networks, at long time scales. This result suggests that actin-microtubule interactions can allow for multifunctionality of active systems – exhibiting steady-state or active responses dependent on the timescale.

Our results provide novel insights into the role of actin-microtubule interactions in controlling motor-driven cellular processes, and the myriad of ways in which the cytoskeleton can modulate self-organized activity. Further, our active system provides a powerful modular platform for introducing and independently tuning various cytoskeleton components (e.g., filaments, motor proteins, crosslinkers, and environmental conditions) in a systematic manner to directly link network composition with the resulting active dynamics. Finally, the ability to introduce reproducible, self-organized activity into a three-dimensional composite network breaks new ground for designing tunable, self-driven, and self-organized active biomaterials.

## Methods

### Cytoskeleton Protein Preparation

Rabbit skeletal actin, porcine brain tubulin, and rhodamine-labeled tubulin were obtained from Cytoskeleton (AKL99, T240, TL590M), and Alexa-488-labeled actin was obtained from Thermo Fisher Scientific (A12373). Unlabeled actin was reconstituted to 2 mg/ml in 5 mM Tris-HCl (pH 8.0), 0.2 mM CaCl_2_, 0.2 mM ATP, 5% (w/v) sucrose, and 1% (w/v) dextran. Labeled actin was reconstituted to 1.5 mg/ml in 5 mM Tris pH 8.1, 0.2 mM CaCl_2_, 0.2 mM dithiothreitol (DTT), 0.2 mM ATP, and 10% (w/v) sucrose. Unlabeled and labeled tubulin were reconstituted to 5 mg/ml with 80 mM PIPES (pH 6.9), 2 mM MgCl_2_, 0.5 mM EGTA, and 1 mM GTP. All cytoskeleton proteins were flash frozen in aliquots and stored at −80°C.

### Myosin Preparation

Rabbit skeletal myosin II (Cytoskeleton, #MY02) was reconstituted to 10 mg/ml in 25 mM PIPES-NaOH (pH 7.0), 1.25 M KCl, 2.5% sucrose, 0.5% dextran, and 1 mM DTT, then flash frozen in aliquots and stored at −80°C. To remove enzymatically dead myosin, 16.7 μM actin was polymerized on ice for 1 hr in 550 mM KCl, 10 mM Imidazole (pH 7), 2 mM Tris (pH 8), 1 mM MgCl_2_, 1 mM EGTA, 0.5 mM DTT, 0.4 mM ATP, 0.1 mM CaCl_2_ and 8.3 μM phalloidin. After polymerization, 1 mM ATP and 2.8 μM myosin II were added to the actin, and the entire suspension was centrifuged at 4°C for 30 min at 32000 rpm. Immediately after centrifuging, the supernatant was removed and stored on ice. Because enzymatically dead myosin binds to actin filaments but does not unbind, the supernatant of the centrifuged solution should contain primarily active myosin.

### Cytoskeleton Network Assembly

Active actin-microtubule networks were polymerized from 2.9 μM each of actin monomers and tubulin dimers in PEM-100 buffer supplemented with 0.1% Tween, 1 mM ATP, 1 mM GTP, 2.9 μM phalloidin and 5 μM Taxol^40^. To image actin and microtubules, 18% of actin monomers and 10% of tubulin dimers were labeled with Alexa-488 and rhodamine, respectively. After the networks polymerized for 1 hr at 37°C, an oxygen scavenging system (45 μg/ml glucose, 0.005% β-mercaptoethanol, 43 μg/ml glucose oxidase, and 7 μg/ml catalase) was added to reduce photobleaching during fluorescence imaging, 0.24 μM myosin was added to introduce network activity, and 50 μM blebbistatin was added to inhibit myosin activity until imaging began.

### Sample Preparation

All samples were loaded into a thin rectangular chamber, formed by adhering a silanized glass coverslip to a glass slide using two strips of heated parafilm (~70 μM thickness). The chambers were immediately sealed with epoxy then imaged. To prevent myosin from adhering to the surface, coverslips were passivated with silane as follows. Coverslips were first plasma cleaned for 30 min, then immersed in a succession of acetone, ethanol, 0.1M KOH, and water baths. Once completely dry, coverslips were immersed in 2% dimethyldichlorosilane, successively immersed in ethanol and water to remove the excess silane, then dried before use. Coverslips were used within a month of silanization^41^.

### Confocal Microscopy

A Nikon A1R laser scanning confocal microscope was used to image networks with a 60× 1.4 NA objective (Nikon) and multiple laser lines and filters to allow for two-color imaging. Microtubules were imaged using a 561 nm laser, and actin was imaged using a 488 nm laser. The 488 nm light also served to inactivate the blebbistatin and initiate myosin activity. Time-series were taken in the middle of the ~70 μm thick chamber for 6 min at a frame rate of 2.78 fps. Examples of time-series are shown in SI Videos 1-3, and representative 256×256 pixel images (212 μm ×212 μm) are shown in Figure 1.

### Particle Image Velocimetry (PIV)

To determine the direction of motion of the filaments during activity, Particle Image Velocimetry (PIV) analysis was performed using the PIV ImageJ plugin^42^. This analysis iteratively measured the cross correlation in image intensity between regions of two images in a time series spaced a lag time *Δt* apart. We chose to perform PIV analysis during four equally spaced time points spanning the 6 min time-series. Each image was divided into 64 regions, from which we generated a field of 64 vectors, each representing the optic flow of a specified region. We chose *Δt* = 20 s for active actin-microtubule networks and *Δt* = 3 s for active actin networks, because the actin-microtubule and actin networks moved a similar distance during these time differences, and motion due to thermal noise was negligible.

### Differential Dynamic Microscopy (DDM)

We performed Differential Dynamic Microscopy (DDM) analysis on each time-series using custom Python scripts^43^. DDM analysis determines how quickly density fluctuations decay by taking Fourier transforms of the difference between images separated by a given lag time *Δt* ^44^. The average Fourier transform squared of all image differences for a given lag time yields the image structure function *D*(*q*, *Δt*). For systems that exhibit a plateau at long lag times, we fit *D*(*q*, *Δt*) to the following model:

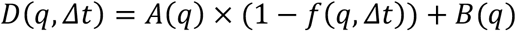

where, *q* is the wave vector, *A*(*q*) is the amplitude, *B*(*q*) is the background, and *f*(*q*, *Δt*) is the intermediate scattering function, which contains the dynamics of the system.

To determine the type of motion and the corresponding rate, we model each intermediate scattering function as an exponential:

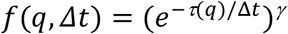

where *τ* is the decay time and *γ* is the scaling factor. We evaluate the decay time *τ* as a function of *q* to characterize the type of motion. For example, if *τ* ~1/*kq* then the system exhibits ballistic motion and *k* represents the corresponding velocity. If *τ* ~1/*kq*^2^, then the system exhibits diffusive motion and *k* represents the diffusion coefficient. The factor *γ* stretches or compresses the exponential and typical values are shown in SI Figure 1. *γ* < 1 characterizes a stretched exponential, while *γ* > 1 characterizes a compressed exponential. Compressed exponentials are indicative of systems undergoing slow ballistic relaxation of internal dipole stresses over larger length scales^45^, which is consistent with stresses generated by myosin motor activity.

### Particle Tracking

As a complementary method for observing active network dynamics, 1 μm polystyrene microspheres were embedded in networks and tracked using custom Python scripts^46^. The beads were labeled with Alexa488-BSA to visualize and prevent non-specific binding to the proteins or surfaces. Time-series of beads were collected at 2.78 fps for 6 min using an Olympus IX73 inverted fluorescence microscope with a 20× 0.4 NA objective and a Hamamatsu ORCA-Flash 2.8 CMOS camera (180 nm/pixel). The bead concentration was such that ~50 beads were tracked in each 1920×1440 pixel field-of-view (346 μm × 259 μm). We use custom Python scripts to measure the frame-to-frame *x* and *y* displacements of the beads and compute the mean-squared displacement (MSD) of the ensemble of beads for lag times of 0.33 s < *Δt* < 343.3 s.

## Acknowledgements

This research was funded by a William M. Keck Foundation Research Grant (awarded to R.M.R.-A., J.L.R., M.D., and M.J.R.). We thank S. Ricketts and B. Gurmessa for work in optimizing and characterizing actin-microtubule polymerization protocols, J. Garamella for help analyzing data, G. Leech for helpful discussion, L. Farhadi for sharing expertise on active actin-microtubule networks, S. Sahu for sharing expertise on coverslip passivation, and V. Yadav and M. Murrell for sharing expertise on myosin II.

## Author Contributions

R.M.R.-A. conceived the project, guided experiments, interpreted data, and wrote the manuscript. G.L. performed experiments, analyzed data, and wrote the manuscript. J.L.R., R.J.M., M.D., and M.J.R guided experiments, interpreted data, and provided useful feedback.

