## supplemental info for "Myosin-driven actin-microtubule networks exhibit self-organized contractile dynamics"

**Supplementary Information**

Videos 1-6 can be found here:

https://drive.google.com/open?id=12dp1_YHhxotztU4ez5Iv4pqDeDIQHw7s

**Video 1. Active myosin-driven actin-microtubule network**

A composite actin-microtubule network contracts due to myosin activity. Actin filaments are green and microtubules are magenta. Both filaments form a well-connected network that contracts smoothly towards the center of the activated region.

**Video 2. Actin channel of the actin-microtubule network shown in Video 1.**

Actin filaments form a well-connected network that contracts smoothly towards the center of the activated region.

**Video 3. Microtubule channel of the actin-microtubule network shown in Video 1.**

Microtubules form a well-connected network that contracts smoothly towards the center of the activated region.

**Video 4. Active myosin-driven actin network**

An actin network contracts due to myosin activity. Actin filaments contract quickly towards local foci that also move quickly and irregularly, causing turbulent dynamics and rupturing of the network.

**Video 5. Inactive actin-microtubule network (+myosin, -UV exposure)**

A composite actin-microtubule network prepared identically to ones shown in Videos 1-3, imaged with no activation of myosin activity (i.e. no blebbistatin inactivation). Only the microtubule channel is shown since the wavelength used to image the actin also de-activates the blebbistatin. The network shows no contractile motion, confirming that the dynamics we measure in myosin-driven networks is a result of myosin activity.

**Video 6. Inactive actin-microtubule network (-myosin, +UV exposure)**

A composite actin-microtubule network prepared and imaged identically to ones shown in Videos 1-3, except for a lack of myosin. The network shows no contractile motion, confirming that the dynamics we measure in myosin-driven networks is a result of myosin activity.

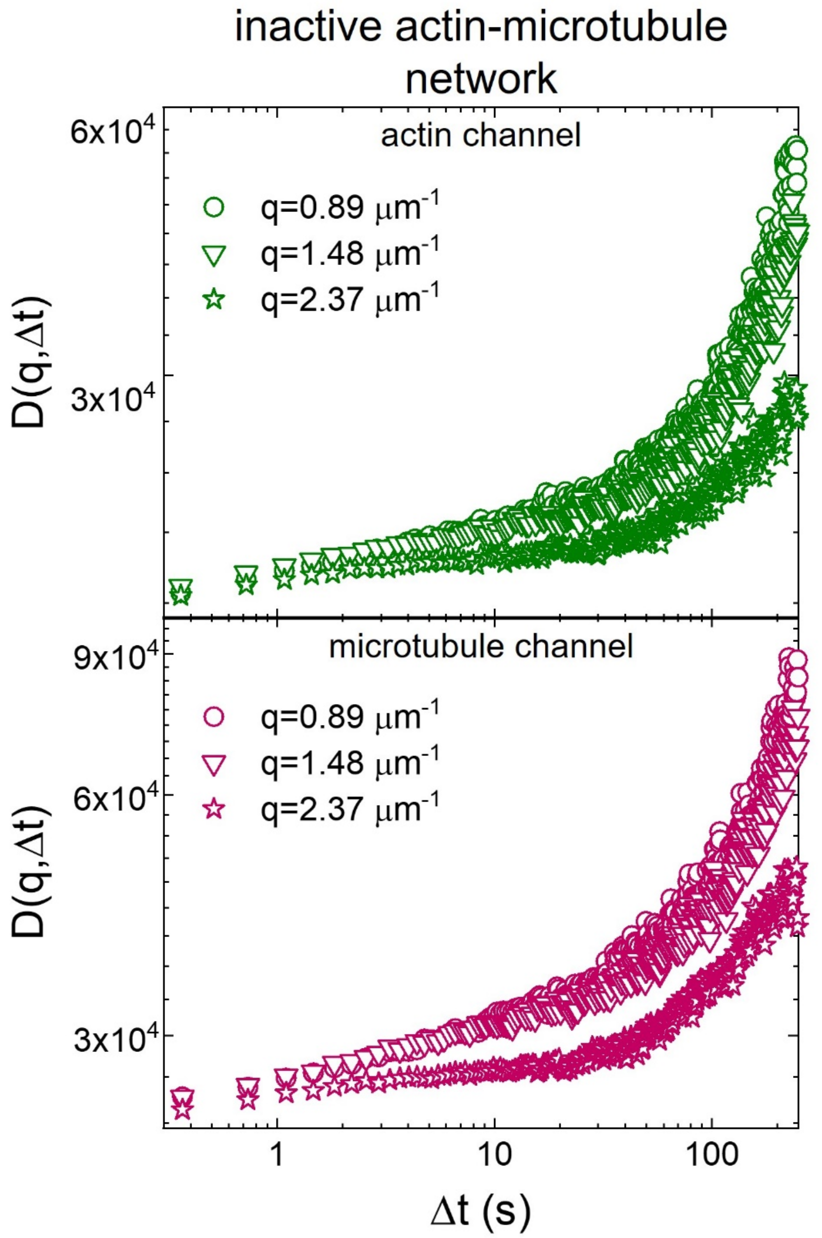


**Figure 1. DDM analysis verifies lack of contractile dynamics in inactive actin-microtubule networks**

Representative image structure functions $D(q,\Delta t)$ for the actin channel (green, top) and microtubule channel (red, bottom) of an actin-microtubule network that contained no myosin, for wavenumbers of $q$= 0.89 $\mu m^{-1}$ (squares, highest curves), 1.48 $\mu m^{-1}$ (triangles, middle curves) and 2.37 $\mu m^{-1}$ (stars, lowest curves). The lack of a plateau for the inactive network demonstrates that the network is nearly static on the timescale of the experiment, compared to the active network, in which both actin and microtubules reach decorrelation plateaus (see Fig 3A).

| network | $\gamma$ |
|---|---|
| active actin-microtubule network (actin channel) | 1.79 ± 0.06 |
| active actin-microtubule network (microtubule channel) | 1.79 ± 0.04 |
| active actin network | 1.46 ± 0.13 |

**Table 1. Active networks undergo slow ballistic relaxation, as characterized by the compressed exponential behavior of intermediate scattering functions.** $\gamma$ factors for intermediate scattering functions (see Methods) of active actin-microtubule networks (top two rows) and active actin networks (bottom row) are shown. Compressed exponentials are characterized by $\gamma > 1$ for all active networks, suggestive of ballistic relaxation of internal dipole stresses generated by myosin motor activity. Both channels of the actin-microtubule network are characterized by identical $\gamma$ factors that are larger and have a smaller spread compared to actin networks.
